# Song complexity in suboscine birds: evolutionary drivers and ecological constraints

**DOI:** 10.1101/2025.10.15.682597

**Authors:** Jingyi Yang, Chiti Arvind, Robert A. Barber, Oscar Johnson, Katie O’Brien, Richard Stanley, Gustavo A. Bravo, Evan J. Buck, Santiago Claramunt, Robb T. Brumfield, Michael G. Harvey, Elizabeth P. Derryberry, Joseph A. Tobias

## Abstract

Acoustic signal complexity varies widely in the animal kingdom for reasons that remain unclear. In birds, it is widely proposed that vocal complexity evolves as an honest signal of individual quality driven by sexual selection. Other hypotheses related to social interactions include competition for ecological resources (social selection) and intra-group communication in group-living animals, both of which may favour signal complexity. However, these hypotheses are rarely explored at macroevolutionary scales, particularly in the context of constraints on sound production, transmission and detection, leading to ongoing uncertainty about the evolutionary origins of complex vocal signals. Using Bayesian phylogenetic models, we test whether different forms of social communication and ecological constraints predict the temporal and spectral complexity of songs in 1,288 species of suboscine passerine birds. We found that song complexity was reduced by sexual selection, along with other limiting factors including large body size and dense vegetation. Conversely, territoriality boosted the temporal complexity of songs. These findings challenge the common assumption that sexual selection is the main driver of increased signal complexity, and instead highlight the role of social selection as a key component of multiple inter-related drivers and constraints.

## Introduction

From the croak of a frog to the intricate serenade of a nightingale, acoustic signals vary widely in complexity across the animal kingdom. Why such variation exists in the apparent sophistication of acoustic signalling systems is a question that has long fascinated biologists [1,2], in part because the answer may help to clarify initial steps along the evolutionary pathway from simple ancestral signals to complex communication systems, including human language [3,4]. However, previous studies attempting to identify the main mechanisms driving the evolution of complex signals have yielded a wide range of conflicting results [5–8] and the relative importance of different hypotheses remains unresolved [9,10] (Figure 1).

**Figure 1.**
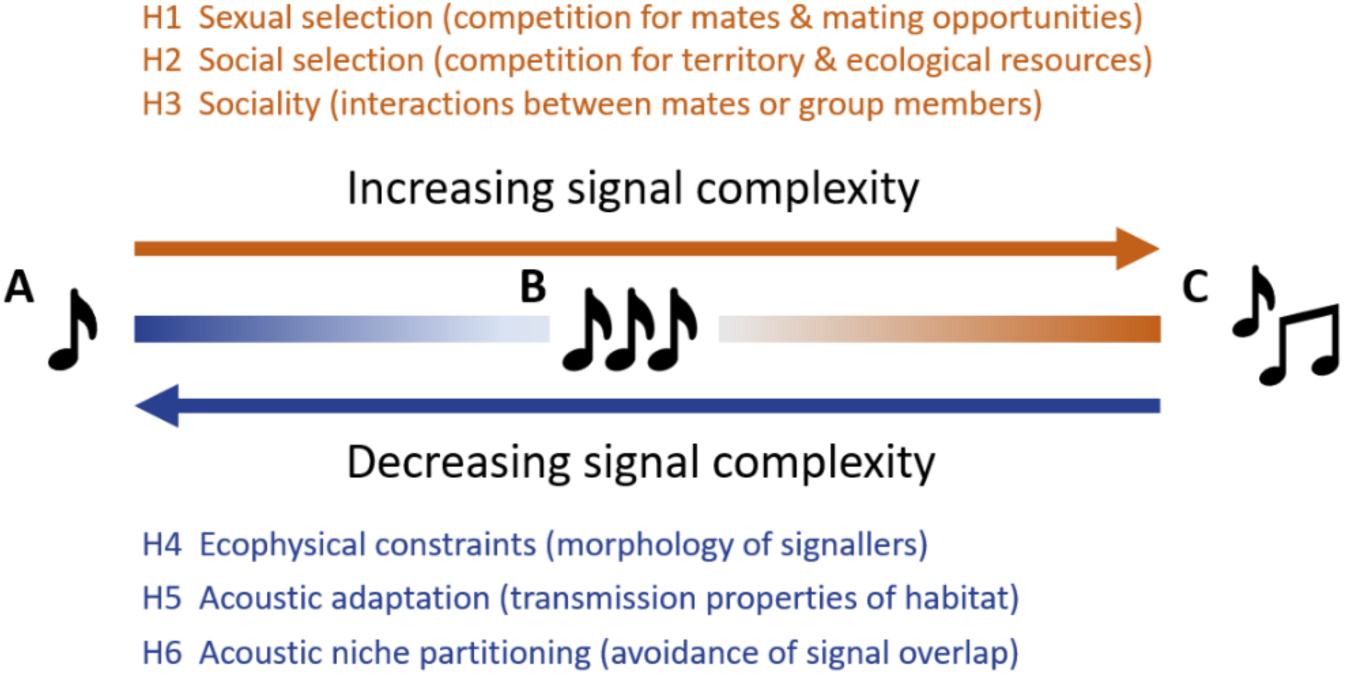
Hypothetical drivers and constraints shaping the evolution of acoustic signal complexity. Using birdsong as a model system, a simplified conceptual gradient of vocal complexity spans from a single note (a) to higher levels of temporal complexity, represented by a repetition of simple notes (b), and ultimately to higher levels of spectral complexity such as multiple note types and higher frequency variation among notes (c). Hypotheses 1–3 (coloured red) are socially-mediated evolutionary mechanisms proposed to drive increased vocal complexity. Sexual and social selection are both proposed to favour more complex birdsongs as honest condition-dependent signals of individual quality used in competition for mates or ecological resources. Signal complexity may also evolve in group-living species (H3) as a result of more complex social interactions and within-group communication. Conversely, hypotheses 4–6 (coloured blue) are physical or ecological constraints limiting or reversing the evolution of signal complexity. According to these constraints, song complexity is disfavoured in species with larger body size (H4), living in dense habitats with poor signal transmission properties (H5), or in species-rich environments where many species compete for the same signal space (H6).

One prominent idea is that complex acoustic signals are costly to produce and therefore function as honest condition-dependent traits indicating individual quality [11–13]. The fact that so many bird species use elaborate songs to advertise for mates in the breeding season and to compete with rivals for mating opportunities suggests that complex signals are the product of sexual selection [14–16]. It seems highly likely, for example, that sexual selection explains the spectacular displays of oscine passerine songbirds with open-ended learning and extensive vocal repertoires, such as mockingbirds [17], starlings [18] and lyrebirds [19]. Based on these observations, birdsong complexity previously served as a standard proxy of sexual selection used in macroevolutionary or behavioural studies [20]. However, some lekking bird species with highly polygamous mating systems produce simple or one-note signals (e.g. cocks-of-the-rock and bellbirds [21]), suggesting that the effect of sexual selection on acoustic signal evolution is relatively unpredictable [8,22]. Recent broad-scale studies seem to confirm that sexual selection is not a consistent driver of high complexity signals, revealing generally weak or reversed associations between polygamy and vocal complexity in birds [9,23].

Two other socially mediated hypotheses have been proposed to explain variation in signal complexity. The first is that signal elaboration, including complexity, may instead be driven by social selection [24,25], with complex vocal signals functioning as acoustic badges of social dominance in the context of resource defence and ecological competition [26]. However, this hypothesis is rarely tested other than through simple scores of the presence or absence of ornamental traits (e.g. [27]), and the extent to which social selection can drive the evolution of acoustic signal complexity is unclear. A third possibility is that sociality itself drives the evolution of signal complexity. For instance, given the varied role of acoustic signals in intra-group communication, complex signals may evolve in group-living animals due to the increased frequency and complexity of interactions between individuals [28–30].

While multiple social mechanisms may select for more complex signals, any increase in complexity is also subject to an array of ecological constraints linked to the physical principles of sound production [31,32], transmission [33], and detection [34,35]. Larger animals tend to produce low-pitched and slow-paced sounds due to biomechanical constraints on sound production organs [36]. In birds, larger beak sizes can place further physical limitations on the pace and complexity of vocal signals [37,38]. In addition, complex or high-frequency sounds can be degraded in dense vegetation [39], driving selection for acoustic signals with slower pace and lower frequency to maximise fidelity of transmission (acoustic adaptation hypothesis) [33,40,41]. Finally, signal detection can be less effective when co-occurring species use signals with similar frequencies [35,42], suggesting that species communicating vocally in environments with high species richness may be restricted to narrower bandwidths or signal space to improve signal transmission and detection (acoustic niche partitioning) [43,44].

Despite the array of potential mechanisms influencing acoustic signal complexity (Figure 1), previous studies have generally tested different social or ecological hypotheses in isolation (e.g. [6,23,30,32]), often focusing on the role of sexual selection [11,16,22,23].

Addressing these hypotheses in conjunction is important given the potential covariance among several predictors such as sexual selection, territoriality, social monogamy and sociality [45,46]. In addition, previous tests have generally been limited to specific clades or regions, typically focusing on relatively small numbers of oscine passerines in the temperate zone [47–49]. These taxonomic and geographic biases lead to suboptimal sampling regimes, including undersampling of tropical systems associated with high levels of social selection [45,50], and patterns of song variation confounded by cultural evolution [51]. Specifically, the songs of oscine passerines are learned and therefore highly plastic, producing extensive variation temporally and spatially (e.g. dialects), as well as within and between individuals (e.g. repertoires), making oscine song complexity highly nuanced [52,53] and difficult to measure robustly [54]. Very few attempts have been made to address these questions at wider geographical and taxonomic scales (e.g. [9]), all of which are based on relatively coarse acoustic characters and patchy phylogenetic sampling.

In this study, we revisit the question of signal complexity by focusing on the songs of suboscine passerines, a diverse radiation of predominantly tropical birds including pittas, broadbills, antbirds, tyrant-flycatchers, ovenbirds and woodcreepers. Specifically, we measured nine acoustic characters representing the temporal and spectral complexity of suboscine songs (Figure 2; Table S1), then used Bayesian phylogenetic models to test six key hypotheses proposed to explain variation in complexity (Figure 1). To do this, we compiled datasets on behaviour, morphology and ecology from literature and assessed the relative contribution of different evolutionary drivers using multi-predictor analyses. This general synthesis is timely because of the recent publication of species-level data on avian morphology and ecology [55].

**Figure 2.**
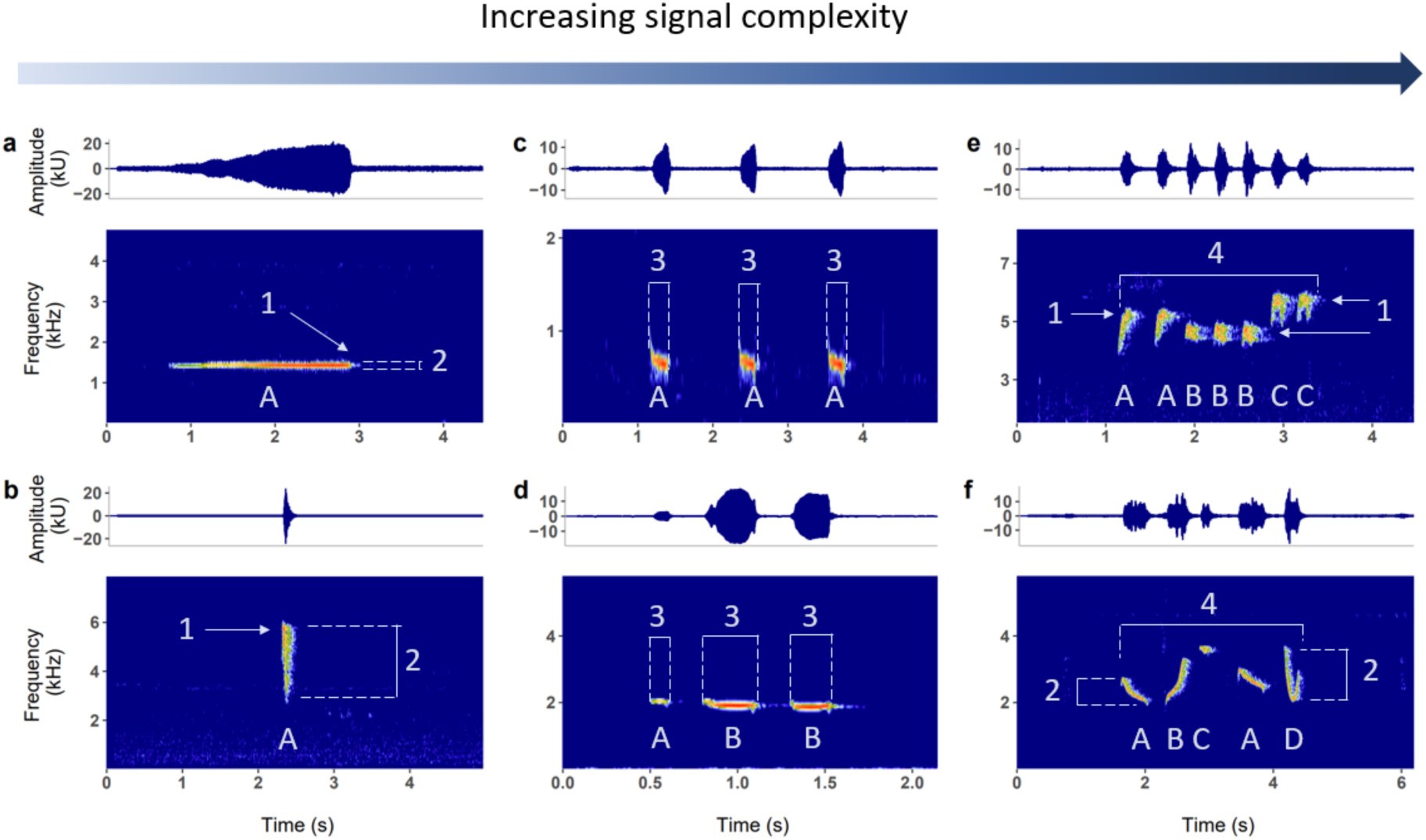
Quantifying acoustic signal complexity from note measurements in songs. Suboscine songs vary from single notes (a, b) to multiple notes (c-f) with varying degrees of temporal (c, d) and spectral (e, f) complexity. For each of 6,465 songs sampled in our study (5.15 ± 1.90 per species), we collected data on eight measurements: note count (the total number of notes), the note type (labelled here as ‘A’, ‘B’, ‘C’, ‘D’), peak frequency (1), bandwidth (2), note length (3), and note slope ( = bandwidth/note length) of each individual note, as well as song duration (4) and note rate (= note count/song duration; Table S1). To quantify song complexity, we used note count, number of unique note types, and overall frequency variation as three main metrics. Frequency variation was calculated as the sum of scaled standard deviations of note peak frequency, bandwidth, and note slope (see Methods). We also used six alternative complexity metrics: song duration, note rate and the standard deviations of note peak frequency, bandwidth, note slope and note length. Complexity metrics based on standard deviations (Table S1) were assigned zero for single-note songs. All complexity metrics were assigned zero for species that do not sing (N = 25) to represent lowest complexity. Exemplar species in spectrograms (a-f): *Erythropitta arquata*, *Casiornis fuscus*, *Hylopezus berlepschi*, *Grallaria flavotincta*, *Sipia nigricauda*, *Schiffornis major*.

Moreover, our analyses also benefit from newly available scores of avian sexual selection [50], in place of standard indices such as size dimorphism and plumage dichromatism, both of which produce misleading estimates of the intensity of sexual selection in passerine birds [56].

Suboscine passerines provide a useful system for understanding signal evolution for several reasons [57]. First, their primary vocal signals are largely innate and stereotyped [58,59], with relatively simple elements showing little variation between song bouts or different individuals, unlike species that learn songs through cultural transmission. The structure of suboscine songs is therefore relatively easy to measure and analyse because they lack features such as unlimited note variation, extensive song repertoires and dialects associated with oscine passerine songbirds ([60]; see Methods). Second, suboscines allow a more rigorous test of social selection and social interactions because the majority of most suboscine families inhabit the tropics where they defend year-round territories or live in groups, unlike most temperate zone songbird species previously studied [50]. Third, the suboscine clade contains over 1200 species with a wide range of behavioural and ecological traits. Their breeding systems vary from strict social monogamy to extreme polygamy, and they occur in a full range of signalling environments from rocky coasts and arid deserts to dense tropical rainforests [57]. Finally, the published suboscine tree [61] is one of the most richly sampled species-level phylogenies available for any major clade, offering an ideal template for exploring variation in signal complexity in the context of evolutionary relationships.

By quantifying how suboscine songs vary in frequency, pace, and syllable composition, and relating this variation to species behaviour, morphology and ecology, we are able to assess the relative importance of different hypotheses proposed to explain the evolution of signal complexity. Our analyses provide a holistic test of three key socially mediated mechanisms – sexual selection [13], social selection [24] and sociality [30] – in the context of widely proposed ecological constraints including morphology [32], habitat density [39], and acoustic niche partitioning among co-occurring species [35].

## Methods

### Study system

We sampled 6,465 songs from 1,276 species (>99%) of suboscine passerines. We define songs broadly as any vocal signals for long-distance communication that typically function in the context of mate acquisition, territory defence or group coordination. Although the term ‘songbirds’ is used exclusively for oscine passerines, the songs of suboscine passerines appear to have identical functions [62]. In most suboscine species, the components of a song (hereafter ‘notes’) are given in a repetitive series or a stereotyped order. Therefore, much of the variation in the vocal complexity of suboscines can be captured by measuring the extent of variation within a single song. We use terms such as ‘song complexity’ or ‘vocal complexity’ of suboscines to refer to the note variation in songs.

### Song selection

Song recordings were sourced primarily from the online acoustic archive Xeno-Canto (xeno-canto.org; n = 5,594), as well as from Macaulay Library (macaulaylibrary.org) and commercial recordings, with a few gaps filled from private collections. When selecting the recordings, we first carefully assessed species identity (see electronic supplementary material). Once identification was confirmed, we measured the most typical advertising songs of each species, with five replicates per species where possible. We selected male songs when these differed from female songs. Both sexes produce songs in many suboscine species, including those with duets and communal vocalisations [27,45]. In the relatively restricted subset of species with sexually dimorphic songs [62,63], we focused exclusively on male songs to avoid pooling data across divergent song types or different selective pressures [62,63]. In species with sexually monomorphic songs, we may have sampled some female songs, but this is unlikely to have affected our analyses because songs had the same (or very similar) acoustic structure and complexity in both sexes.

In a few suboscine species that do not have songs yet habitually vocalise, we used the most complex long-range calls (i.e., the loudest call type with consistent tone and structure) that are likely used in long-distance intraspecific communication. These intraspecific calls were only used when the species is not known to have more complex vocalisations based on literature information and field experience (e.g. *Xipholena punicea*). We also carefully excluded atypical songs (e.g. considerably faster or longer than usual after being agitated by playbacks), alarm calls (e.g. abrupt with unusually high bandwidth), and non-vocal acoustic signals (e.g. wing snapping of manakins). In rare cases where there was no recording available, we scored the species as either unknown or as having no primary vocalisation (complexity of zero). We determined this score based on the rarity of the species and the vocal behaviour of its closest relatives (see electronic supplementary material).

For species that are known to sing or produce long-range vocal signals (n = 1,251 species), we selected the best quality unaltered recordings and the clearest song within that recording for analysis. Good quality recordings were defined as those with no overlapping vocalisations, high signal-to-noise ratio and low reverberation. Occasionally, we selected two songs per recording if there were limited recordings for the species, in which case we chose the highest quality recordings and analysed two well-separated songs within each recording. Our final dataset contained 6,465 songs (5.15 ± 1.90 songs measured per species).

### Note measurement

We analysed the selected songs using the software Raven Pro v1.6 (Cornell Lab of Ornithology). To ensure consistency of measurements, we used a set of established spectrogram parameters following Tobias et al. [60] and manually selected all notes of the song (see electronic supplementary material). A note was defined as a continuous trace on the spectrogram and was individually measured from its fundamental frequency. The songs of some species consist of a single note, and if given in sequence, the number of songs and their intervals vary widely (e.g. *Procnias* bellbirds). For those single-note songs, we measured as many high-quality songs in one recording as possible, then averaged the measurements to minimise measuring errors (see electronic supplementary material). In a few species with more than one primary vocalisation, including songs with either single or multiple notes (e.g. *Sirystes subcanescens*), we sampled different song types in the ratio that they appeared across all recordings. In very few cases where multiple different song-types are produced (e.g. *Attila spadiceus*), we selected the most complex phrase to represent the highest vocal complexity. When measuring notes on the spectrogram, we carefully distinguished different singing individuals (e.g. in duetting species such as *Culicivora caudacuta*) and excluded any non-focal species.

### Song complexity metrics

Complexity is difficult to quantify [54,64], so we chose metrics based on different dimensions of complexity. We assume that complex vocal signals consist of a larger number of (1) notes or (2) unique note types, as well as (3) wider variation in the frequency of different notes [9,23].

Accordingly, when analysing songs, we first recorded the total number of notes present in that song (‘note count’) and the number of unique note shapes (‘note type’) by visually inspecting the spectrogram. We then quantified frequency variation by summing the standard deviations of the peak frequency, bandwidth, and slope of all notes in the song (‘frequency variation’), with the input variables first z-standardised to ensure compatibility (Table S1). These three variables were then used as the main metrics of suboscine song complexity. To further quantify the temporal variation of songs, we also considered the standard deviation of note length, song duration and note rate (= note count/song duration), the latter two metrics are also often used to represent the singing effort of the species (Figure 2, Table S1; see electronic supplementary material). For single-note songs, we assigned zeros to all complexity metrics based on standard deviations (see Table S1) to reflect the absence of inter-note variation in those songs, and note rate was calculated as the number of songs given per second to allow comparison with multi-note songs (see electronic supplementary material). For silent species (n = 25), we assigned zeros to all metrics to indicate the lowest complexity. Finally, we calculated the median value of each complexity metric as the final complexity scores for each species.

### Social and ecological predictors

#### Social behaviours

To quantify the intensity of sexual selection in suboscines, we used the five-point scoring system following Barber et al. [50], who classified the degree of polygamy based on the dominant social mating system of the species, adjusted by the prevalence of extra-pair paternity (EPP; Data S1). Sexual selection scores range from 0 (strict monogamy) to 4 (extreme polygamy) to represent the increasing levels of reproductive competition and skewed success among males. The intensity of social selection was quantified as the territoriality of the species, with data adapted from Tobias et al. [45]. We combined seasonal and permanent territoriality as ‘territorial’ (as opposed to non-territorial) in the main phylogenetic models, since we expect signals of social dominance to mediate effective territory defence even in seasonally territorial species. To quantify the degree of sociality we collected data on the maximum social group size. A social group was defined as a stable aggregation of individuals demonstrating strong social cohesion and interaction on a long-term, typically year-round basis, excluding any temporary flocks (see electronic supplementary material). We collected group size data primarily from species accounts in *Birds of the World* [65]. From the text on sociality, breeding and foraging, we estimated the maximum stable social group size for each species (see electronic supplementary material).

Behaviours of some species are poorly known, especially in the tropics [50]. The information for those species was based on incomplete observations and sometimes can only be inferred from their close relatives. To account for the differences in data certainty, each of the social behavioural data (sexual selection, territoriality and social group size) was accompanied by a certainty score which ranged from A (highest certainty) to D (lowest certainty; Data S1) following Tobias et al. [45].

### Ecological conditions

Larger body size has been shown to affect signal design [36]. As a corollary, in birds, larger beaks may further reduce motor performance and frequency variation, placing constraints on signal complexity [32,66]. Therefore, to test the effect of morphological constraint on song complexity, we included body mass and beak size in models with data obtained from AVONET [55]. Body mass was calculated as the midpoint average of all mass data reported for the species. Following Derryberry et al. [40], we calculated beak size as the product of beak length, width and depth to represent the three-dimensional size of the beak. Since absolute beak size is largely determined by body mass, we divided beak size by body mass and used this relative size metric in models to test the independent effect of beak size.

To quantify habitat density, we adapted the habitat data in AVONET [55] by reversing the values so that larger numbers indicate denser habitats, therefore reflecting greater constraints on signal complexity (Data S1). Habitat data were designed to reflect the physical condition of signal transmission. Thus, although dense habitats generally involve forests or tall shrublands while open habitats are grasslands and deserts, exceptions were made depending on the specific behaviour of the species. For instance, species living in forests but habitually singing on top of the canopy were classified as open-habitat dwellers [45]. Conversely, species that live in swamps or wetlands were classified as dense-habitat dwellers if they tend to sing low in dense vegetation such as reedbeds (Data S1). Finally, to test the hypothesis of acoustic niche partitioning, we used species richness data obtained from Harvey et al. [61], who estimated the total number of suboscine species with >1% of geographical range overlap with the focal species in their native and breeding distributions (Data S1).

### Phylogenetic modelling

All analyses were conducted in R version 4.2.2 [67].

To test how song complexity covaries with social and ecological conditions, we ran Bayesian phylogenetic generalised linear models using the R package *brms* v2.18 [68]. The response variable was one of the nine complexity metrics (Table S1). Species that are known not to sing (n = 25) were also included in models with all complexity metrics assigned zero, while unknown species (n = 12) were removed from analyses, resulting in a final sample size of 1,276 species.

Given the different distribution types of the response variables, we used different families when fitting *brms* models. For count data (note type and note count), we first rounded data to integers then fitted with negative binomial family; for continuous data (all other metrics), we first normalised the raw data with square root transformation then used the Gaussian family.

In each model, we included all seven social and ecological factors as explanatory variables, and used the newly assembled phylogeny of all suboscine species [61] to account for the phylogenetic non-independence between species [69]. When modelling complexity metrics derived from standard deviations (i.e., frequency variation, and the standard deviations of peak frequency, bandwidth, and note slope), we also added mean peak frequency (Hz) as an extra explanatory variable, since the complexity score of high-pitched songs can be systematically high for mathematical reasons (see electronic supplementary material). Similarly, we added mean note length (s) as an extra explanatory variable to the model of note length variation. To improve normality, mean peak frequency and note length were square-root transformed while all other continuous predictors were log-transformed. All continuous predictors were then scaled to a standard deviation of 0.5 to facilitate comparison of effect sizes between continuous and categorical variables [70]. Semi-continuous variables (sexual selection scores, habitat density) were untransformed. Given the different taxonomies involved in the source datasets, we aligned all original data to the Clements checklist v.2017 [71] used in the suboscine phylogeny [61]. Species lacking data due to taxonomic mismatches were given new data that were either generated using the original methods (i.e., species richness), or transferred from the nominate species in the original dataset (i.e., morphological traits; see electronic supplementary material).

Following standard procedures [72,73], we used weakly informed Bayesian priors in all models: *Normal(0,1)* for the estimated slopes of all predictors and *Normal(0,2)* for intercepts.

Each model was run with 15,000 iterations (the first 5,000 iterations were discarded as the burn-in) using 4 chains and a thinning rate of 20, leading to the final effective sample size of 2,000. Convergence of chains was inspected visually and with the r-hat value reaching one (Table S2). There was no collinearity found between explanatory variables (variance inflation factors: 1.01-1.19; see also Figure S1), and we report statistical significance as the 95 credible intervals (CI) do not span zero. Due to higher uncertainty in social behaviour data, we repeated all models using high certainty data only (i.e., certainty = A and B; n = 1,028 species). The results for all species and the restricted sample of higher-certainty data were nearly identical (Table S2), so for clarity we only report model results based on data of all species.

## Results

Acoustic signal complexity varied widely in suboscines from single-note songs (n = 148) to songs with long duration (e.g. *Scytalopus opacus*, median note count = 432) or high spectral complexity (e.g. *Empidonax oberholseri*, median number of note types = 9). A minority of species (n = 25; 1.9%) were deemed songless because they seemed to produce no long-distance vocal signal. Song complexity also seemed to show phylogenetic patterns, with the highest note counts found among tapaculos (Rhinocryptidae) and woodcreepers (Furnariidae), and the most note types among tyrant flycatchers (Tyrannidae; Figure 3). Contrary to predictions, we noticed that polygynous clades (e.g. Cotingidae) have lower average number of note types and contain most silent species (Figure 3a). In contrast, songs of territorial clades (e.g. Rhinocryptidae) tend to show substantially higher note counts (Figure 3b).

**Figure 3.**
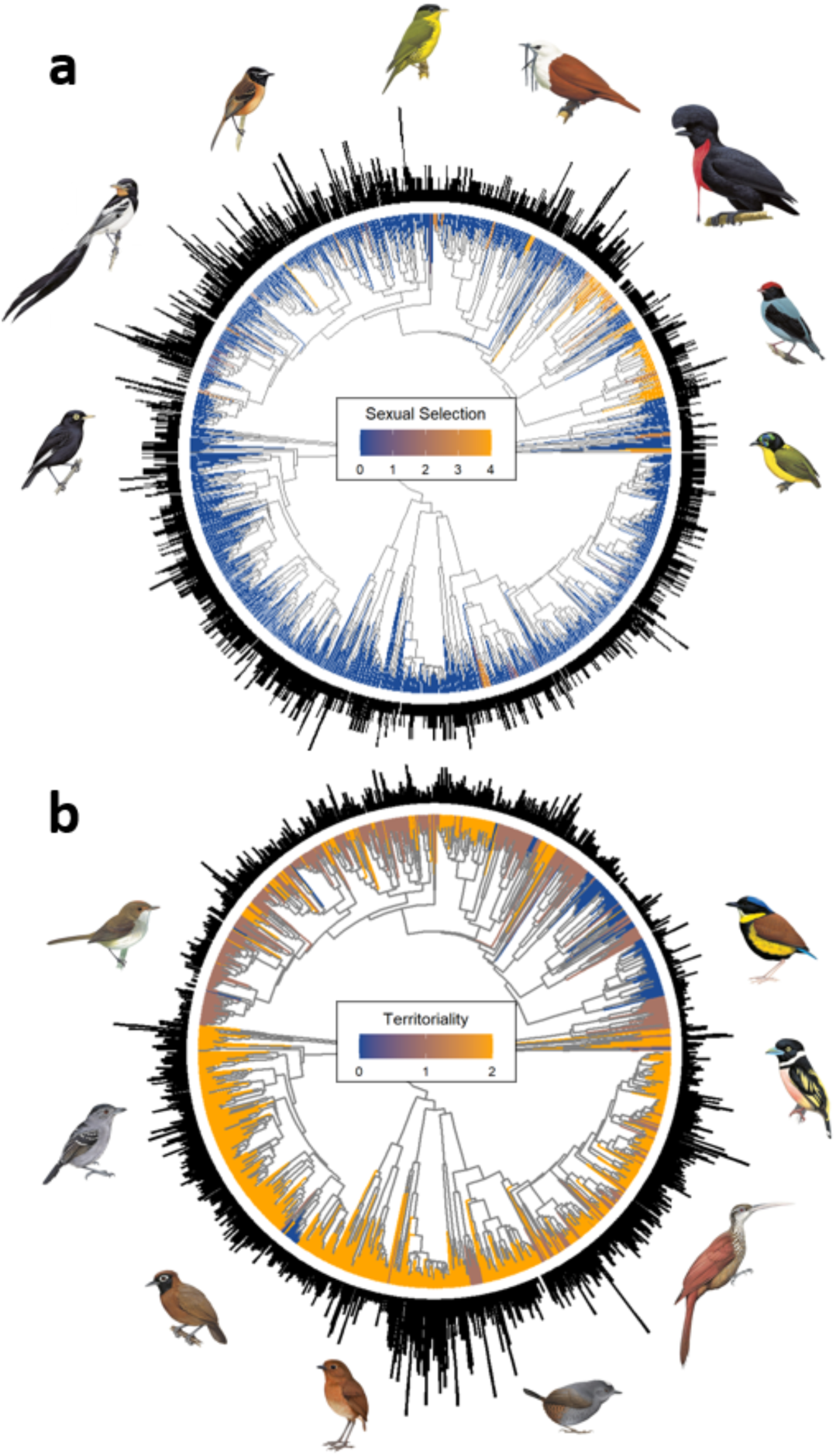
Song complexity in suboscine passerine birds (N = 1,276 species). Phylograms show how spectral and temporal complexity varies across the suboscine radiation (n = 1,276 species). For each species, bar length at the tip of branches represents (a) the number of unique note types, and (b) note count. On branches, warmer colours represent higher estimated intensity of (a) sexual selection and (b) territoriality (as a metric of social selection). Sexual selection scores are from 0 (strict monogamy) to 4 (extreme polygamy), and territoriality scores are from 0 (non-territorial) to 2 (year-round territory defence), based on published species-level datasets [45,50]. Note count data were square-root transformed for visualisation. Bird images are reproduced with permission from Birds of the World (Cornell Lab of Ornithology) and Lynx Nature Books (see acknowledgements for image credits).

The raw patterns were confirmed by the Bayesian phylogenetic model results (Figure 4; Figure S2; Table S2). Sexual selection scores were significantly related to lower number of note types (CI = [-0.15, -0.01]), while territoriality was associated with higher note counts (CI = [0.03, 1.02]; Figure 4). These results indicate that social selection, rather than sexual selection, better predicts signal complexity in suboscines. Our models also revealed that the effect of sociality was largely absent (Figure 4; Figure S2), suggesting that group living is not associated with signal complexity in this clade.

**Figure 4.**
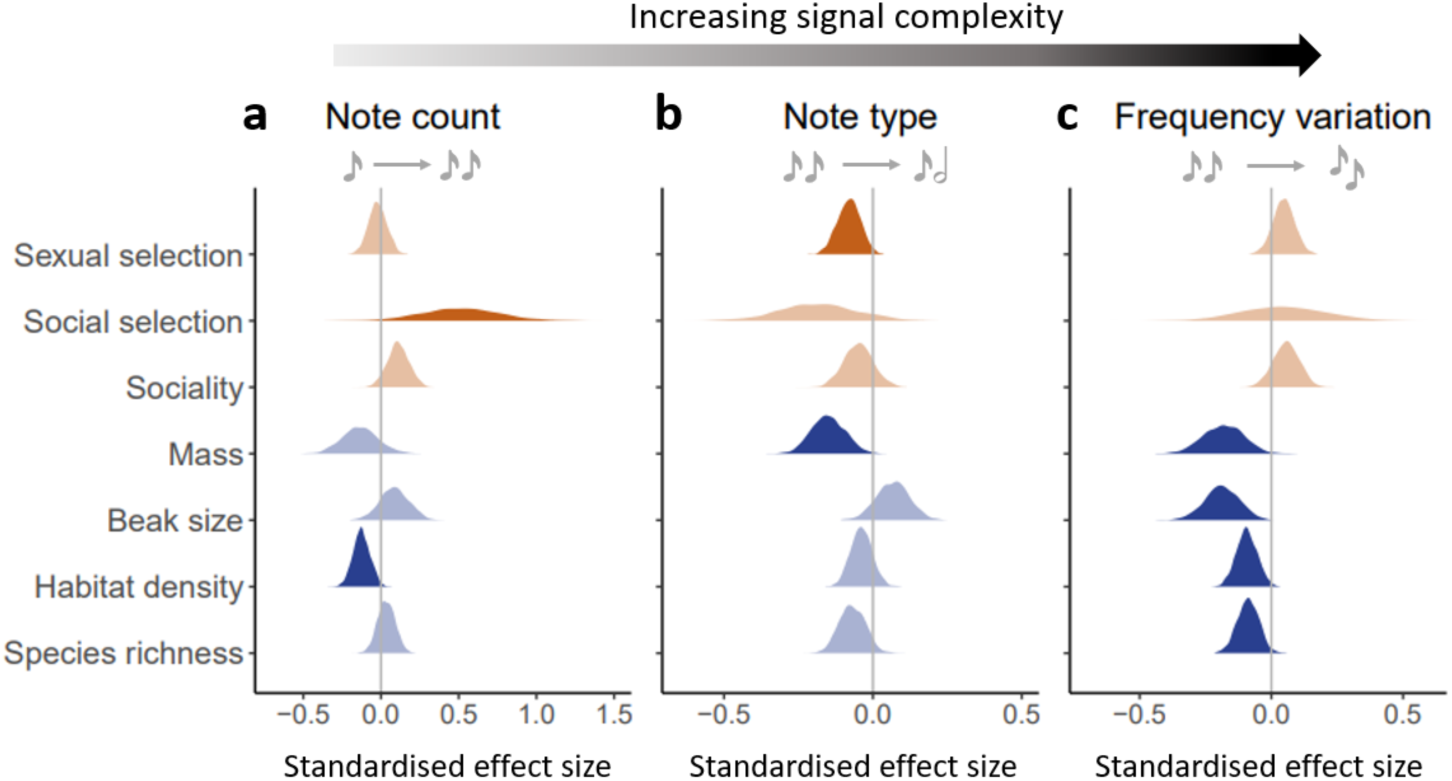
Social and ecological predictors of birdsong complexity. Panels show the results of Bayesian phylogenetic models testing the effects of the proposed social drivers (red) and ecological constraints (blue) on the evolution of song complexity in 1,276 suboscines species. Temporal and spectral complexity was estimated as note count (a), the total number of unique note types (b), and variation in note frequency (c). Frequency variation was measured as the total variance in the peak frequency, bandwidth and note slope of all notes in the song (see Methods). Models used the phylogenetic tree provided in Harvey et al. [61]. Each coloured area shows the distribution of estimated effect sizes of the predictor, based on a sample of 2,000 posterior draws. Dark shades indicate statistical significance where 95% credible intervals do not span zero. To account for the statistical artefact related to complexity metrics using standard deviations (see electronic supplementary material), the frequency variation model (c) included one additional explanatory variable, the average frequency of all notes, which is not shown in the panel for visual clarity. See Table S2 for the full statistics of the models. Repeating the same analyses with alternative complexity metrics (Figure S2) and a conservative sample of high-certainty data (Table S2; see Methods) produced very similar results.

Compared to socially mediated drivers, we found stronger and more consistent support for ecological constraints. For example, large body size was significantly associated with lower note type (CI = [-0.26, -0.04]) and frequency variation (CI = [-0.33, -0.04]; Figure 4), while larger beak size also reduced the latter (CI = [-0.32, -0.06]; Figure 4). The effect of habitat density was similar, with denser habitats significantly associated with lower song complexity including lower note count (CI = [-0.23, -0.02]) and frequency variation (CI = [-0.17, -0.01]; Figure 4). We also detected a weak negative effect of species richness, which was associated with lower frequency variation (CI = [-0.17, -0.01]) but no other complexity metrics (Figure 4).

To inspect signal complexity in greater detail, we decomposed frequency variation into its underlying measurements (variations of peak frequency, bandwidth, and note slope) and used three additional metrics of temporal complexity (variation of note length, song duration and note rate) before repeating the models. We found that the negative effects of body mass and beak size on signal complexity remained (i.e., both factors tended to reduce spectral complexity; Figure S2a-c), while large body mass was additionally associated with higher variation of note length, longer song duration and slower note rate (Figure S2d-f). We also found that denser habitat was significantly associated with lower spectral complexity (Figure S2bc) and lower note rate (Figure S2f), in line with the prediction of the acoustic adaption hypothesis.

## Discussion

Using comprehensive trait and phylogenetic data for suboscine passerine birds, we showed that acoustic signal complexity co-evolves with a range of social and ecological factors, in particular social selection, species morphology and habitat density. Our analyses reveal that song complexity in suboscines is reduced by sexual selection and increased by social selection (Figure 4). We also found consistent effects of ecological constraints on signal complexity, with lower complexity songs generally associated with larger body size and beak size, and occurrence in denser habitats (Figure 4; Figure S2). In contrast, we found evidence that sociality and acoustic signal partitioning in multi-species assemblages have negligible impact on song complexity in this clade. The picture emerging from this diverse pantropical radiation is that the evolutionary origins of signal complexity are highly nuanced and multidimensional, requiring much more detailed exploration of confounding variables than most studies allow for.

### Social drivers

A key finding is that sexual selection has opposite effects on signal complexity than often proposed in studies focusing on animal species from higher latitudes. Although long hypothesised as the key driver of higher complexity in avian acoustic signals [12–14], we found no evidence that sexual selection increases song complexity in any spectral or temporal aspect (Figure 4; Figure S2). An increased strength of sexual selection was not significantly associated with note count or frequency variation, and indeed was negatively associated with the number of note types in the song. This is opposite to the pattern initially described by Darwin [13], therefore rejecting the sexual selection hypothesis. Our finding in suboscines aligns with other recent large-scale analyses on birdsongs [9,23], suggesting that sexual selection is not a general mechanism driving the evolution of song complexity. One reason for this outcome is that acoustic signals may be sexually selected whereas song complexity is not. In some taxa, song performance is thought to be assessed by females via the loudness [21,74], trill rate [75,76] or consistency [12] of songs instead of higher variation. This may also apply to suboscines as the songs of some polygynous species are simple but extremely loud (e.g. *Procnias* bellbirds) or low-pitched (e.g. *Cephalopterus* umbrellabirds) [21]. Overall, the reduced number of note types identified in polygynous lineages suggests that sexual selection does not necessarily promote evolutionarily labile acoustic signals [77] and instead may favour extreme signal simplicity and stereotypy, particularly when songs need to be broadcast and detectable over very large distances to attract mates in low density populations (e.g. cotingas).

Another reason why sexual selection in suboscines may reduce song complexity is the potential trade-off between costly vocal elaboration and other targets of sexual selection. In suboscines, highly sexually selected species are often adorned with elaborate plumage ornaments (Figure 3), courtship displays (e.g. *Chiroxiphia* manakins) and non-vocal sounds [78]. Maintaining multiple elaborate signals may be energetically costly or decrease signal clarity, so species are predicted to have either elaborate songs or plumage, but not both (transfer hypothesis; [79]), and to favour one signal modality rather than multiple modalities (redundancy hypothesis; [80]). Different costly and redundant signals may therefore compete with each other in multi-modal communication systems [80]. Evidence for this trade-off has been reported in parrots [81], hummingbirds [82] and suboscine passerines [83], suggesting that song simplicity may be offset by elaborate plumage in some highly polygynous species [56]. Apparent trade-offs between vocal and visual signals can also be reversed in suboscines, as illustrated by the Screaming Piha (*Lipaugus vociferans*), a polygynous cotingid with a complex song and very dull plumage. Nonetheless, the transfer and redundancy hypotheses have received only patchy support and require further testing in passerine birds.

Few previous studies have systematically assessed the role of social selection in signal evolution [27,84]. Our results show that social selection is likely a key driver of signal complexity, especially along the temporal dimension. Seasonal and year-round territoriality is significantly associated with increased note count (Figure 3; Figure 4) and tends to increase song duration and note rate (Figure S2e-f). We did not detect positive effects of territoriality on spectral variation, perhaps due to constraints on motor performance during fast sound production, making it hard to produce variable frequencies at a high rate [85]. However, it is possible that increased note count represents an initial step towards higher signal elaboration because it allows further spectral modifications to an increased number of components in a signal.

Contrary to predictions, sociality showed no positive effects on song complexity except an increased note slope variation and note rate (Figure 4; Figure S2). This may seem counter- intuitive since social species have been shown to develop larger repertoire sizes [10] and even various call types with different referential functions (e.g. *Psophia* trumpeters; [86]). However, those vocalisations may be fundamentally different from advertising songs in that the former are mainly close-range contact calls designed for within-group communication. The complexity of those vocalisations was not included in our study due to much lower data certainty.

Moreover, a larger repertoire of vocal signals may be functionally complex yet structurally simple. Therefore, while our study shows that group size did not predict the complexity of long-distance vocal signals in suboscines, sociality may still increase signal complexity in other aspects [87].

### Ecological constraints

In contrast to social drivers, we found more consistent effects of ecological factors (Figure 4, Figure S2) especially species morphology. Large-bodied species sing with fewer note types, lower frequency variation (Figure 4) and slower pace (Figure S2), corroborating the role of physiological constraints on sound production. The effect of beak size was similar, although slightly weaker (e.g. no effects on the temporal complexity; Figure 4, Figure S2), potentially because some complex songs can be produced without fast beak movements but instead using specialised syringeal muscles [88]. This alternative may be particularly useful when beak design is simultaneously under selection for multiple conflicting ecological functions including singing, foraging and thermoregulation [89], which in turn can further decouple the relationship between beak size and song complexity.

We found that increased habitat density was associated with lower song complexity both temporally (e.g. note count, note rate) and spectrally (e.g. frequency variation), adding further support for the acoustic adaptation hypothesis [40]. In contrast, the negative effect of species richness was only detectable on frequency variation (including note slope variation) and no other metrics, suggesting that acoustic partitioning is unlikely to be a major constraint on signal complexity. While reduced frequency variation in species with many co-occurring relatives may be a consequence of competition for signal space, this hypothesis requires further testing. For example, it is possible that our analyses do not fully account for the fact that the highest levels of suboscine species co-occurrence occur in tropical forests, a signalling environment which also imposes constraints on complexity because of the denser habitat. This covariance between vegetation and interspecific signalling environments highlights the importance of multi-variate models accounting for correlated effects. In any case, previous research has shown that closely related sympatric suboscine species may sing at similar times and with convergent signal structures [90] and that females are adept at discriminating between species with near-identical male songs [91]. These observations are opposite to the predictions of the acoustic niche partitioning hypothesis, suggesting that species recognition in co-occurring suboscine species is not necessarily a cognitive challenge for receivers.

### Limitations

Species attribution is generally accurate in online sound archives as a result of direct curation by a global community of song identification experts, so misidentifications are unlikely to affect our results. However, other factors may limit the accuracy of song complexity measurements. For example, estimates could be affected by variation in amplitude (which can change song structure by introducing strong harmonics or reverberation), background noise (e.g. conspecific and heterospecific birds, as well as mammals, amphibians, insects), and mistaken classification of vocal types (e.g. mixed samples of songs, flight calls, alarm calls, contact calls). These issues can be particularly problematic in highly diverse tropical environments where vocal signals from multiple individuals or background species in the same recording can change the apparent number of notes and note shapes in a focal song. One example of an intraspecific complication is the highly synchronised display calls of *Chiroxiphia* manakins, in which multiple males produce overlapping acoustic signals, potentially inflating complexity estimates. The difficulty of detecting background noise contributing to apparent song structures may result in high uncertainty in large-scale studies using automated methods to detect songs and analyse their vocal parameters (e.g. [9]). In our study, we largely circumvent these problems by careful checking and curation of samples to discard song recordings with overlapping elements from non-focal individuals or species, and to manually segment standardised elements of songs (see electronic supplementary material).

Although our sampling procedure should provide relatively robust estimates, it is clear that song complexity is inherently difficult to quantify [54]. Since we are using human hearing and on-screen measurements, it could be argued that our approach overlooks the fine details in acoustic signals that are likely to be detected by avian receivers [92]. While sound perception is more refined in birds than humans [91], previous research has shown that humans share remarkable neural similarities with birds related to acoustic signal production and perception [93,94], suggesting that human scores of birdsong structure are likely to correlate with signal complexity perceived by birds [95]. Finally, we have focused on the internal acoustic structure of primary song types rather than overall size of the vocal repertoire, which in birds is usually estimated as the number of different songs, phrases, or other acoustic signals, used by an individual over time [18,86]. Our approach therefore provides an incomplete test of sexual selection and social interactions as drivers of vocal complexity [87]. However, we emphasise that our goal is to test initial steps along the pathway towards signal complexity so the stereotyped acoustic signals of suboscine passerines make an ideal system largely uncoupled from the complexities of vocal repertoires [57].

## Conclusions

Interspecific variation in signal complexity is relevant to several prominent themes in evolutionary biology, including individual recognition [96,97], species interactions [98–100], and speciation [34,101]. In addition, given the convergent neural apparatus for vocal learning in birds and humans [93], birdsong complexity has long been viewed as a model system for understanding linguistic, cognitive and cultural phenomena [94,102]. By focusing on a highly diverse group of species with minimal vocal learning [59,103], we strip away the effects of cultural evolution to address the underlying factors promoting complexity in birdsong. Our results challenge the common assumption that sexual selection is positively related to complexity, and suggest instead that territorial behaviour and ecological constraints play more significant roles in shaping song complexity at macroevolutionary scales. Moreover, we find evidence that sexual selection reduces note variation whereas social selection boosts temporal complexity of the song. These results provide a more unified understanding of acoustic signal evolution, and suggest that birdsong complexity reflects trade-off among multiple social and ecological factors playing out along different temporal and spectral dimensions.

## Acknowledgements

We thank Alex Cranston, Charis Declaudure, Stephanie Kam, Lois Nolan, Harry Peck, Edward Wickstead and Lily Wright for help with data collection. Funding was provided by the Silwood Park masters programme (Imperial College London) and the US National Science Foundation (Award DEB-2203216 to M.G.H.; Award DEB-1146265 to R.T.B.). We thank the many sound recordists who contributed recordings to the databases used in this study. The credits for species images used in Figure 3 are as follows: (panel a) *Hymenops perspicillatus* and *Alectrurus risora* by Hilary Burn; *Polystictus pectoralis* by Brian Small*, Laniisoma buckleyi, Procnias tricarunculatus* and *Cephalopterus glabricollis* by Chris Rose*, Chiroxiphia caudata* by Jan Wilczur*, Philepitta schlegeli* by John Cox; (panel b) *Euscarthmus meloryphus* by Brian Small*, Thamnophilus atrinucha* and *Rhegmatorhina cristata by H*ilary Burn*, Grallaria ayacuchensis* by Liz Wahid*, Scytalopus sanctaemartae* by John Cox*, Nasica longirostris* by Tim Worfolk*, Eurylaimus ochromalus* by Ian Lewington*, Hydrornis gurneyi* by Chris Rose.

## Supplementary Information

### Supplementary methods

#### Data collection

##### Checking species identity

When sampling suboscine species, we followed the taxonomic framework of the Clements Checklist v2017 [1], in alignment with a species-level suboscine phylogeny published by Harvey et al. [2]. Given the constant taxonomic flux and occasional misidentification of species in acoustic archives, we corroborated species identity with published information and acoustic data. First, we compared the content of sound files against song descriptions presented in literature such as *Birds of the World* [3]. We then cross-validated songs by checking other recordings of the same species across different individuals and regions where possible. Almost all suboscine songs in Macaulay Library and the Xeno-canto archive are correctly identified after intensive curation by a global community of bioacoustics experts who routinely report inaccurate data.

### Intraspecific variation

Suboscine songs often vary geographically, linked to underlying genetic variation [4]. In most cases, this intraspecific vocal variation is limited to the pace or rhythm of the song as larger differences are routinely used to determine species limits in suboscines [5]. When sampling songs, we attempted to restrict data collection to similar vocalisations within a particular subspecies or set of subspecies producing similar songs. This was not possible in some species where the total number of available sound recordings was limited, meaning that we were forced to sample across different geographical song types. However, in all such cases the level of song complexity was similar across different song types.

We tried to avoid sampling aggressive songs recorded after playback as these can contain large numbers of notes in some species such as tapaculos (Rhinocryptidae). This was not always possible so some estimates of note number may be inflated by post-playback recordings. However, song complexity in these cases reflects natural variation that occurs during agonistic interactions between territorial individuals.

### Determining ‘songless’ species

When there was no song recording available for a species, we carefully assessed whether the species is silent. That is, if the species is well-studied and had no songs recorded, we treated that species as silent (n = 25 species). If a species potentially sings given many of its close relatives are known to sing, but did not have recordings because of its rarity, we treated that species as unknown to avoid underestimating its signal complexity (n = 12 species; Data S1).

### Note measurement

To ensure consistency of note measurements, we used the following spectrogram parameters in the Raven software: window = Hann, DFT (discrete Fourier transform) size = 1024 samples, hop size = 128 samples, and overlap = 87.5%, as specified in ref. [6] with similar parameters widely used in avian acoustic research [7,8]. We define a note as a continuous trace on the spectrogram. For the majority of songs, we measured all notes of the song individually, including trills, as long as the notes appear as separate traces. Some species produce long songs by repeating the same note for over 50 times (e.g., *Scytalopus* tapaculos). For those simple long songs, note measurements (except note count) were taken from 10 consecutive notes from the beginning, 10 in the middle and 10 at the end. For other long songs consisting of different note types (e.g., *Xiphorhynchus* woodcreepers), we measured all notes individually to ensure the full coverage of note variation. In some species, songs are given as a single note only (e.g., *Cephalopterus* umbrellabirds), and if the notes are given in sequence, the total number and intervals between those notes are highly unpredictable with no clear start and end. Therefore, this song type is fundamentally different from normal multi-note songs, and were treated as single-note songs. For single-note songs, we measured all notes (songs) in the recording and averaged values to minimise measuring errors. Because each song contained only one note, we calculated complexity metrics in a different way (see next section).

In rare cases, the division between single- and multi-note songs is not clear. For example, the dawn song of *Sublegatus arenarum* consists of two simple notes alternating at similar intervals for an unknown period of time. We therefore measured the two notes together, to reflect its higher complexity than single-note songs and lower temporal complexity than longer, regular multi-note songs.

### Complexity metrics

To calculate complexity metrics, we first counted the total number of notes (note count) and the number of unique note types (note type), and exported the following raw measurements for each note from the Raven selection table: begin time (s), end time (s), peak frequency (Hz), bandwidth 90% (Hz). Peak frequency is where the note reaches its highest amplitude, and bandwidth 90% (hereafter ‘bandwidth’) is the difference between the two frequency limits spanning 5% and 95% of the energy of the selected note. We chose these two energy-based frequency measurements since they are less sensitive to different recording conditions and potential measuring errors, and thus provide a more robust estimate of frequencies [8,9]. From these measurements, we then calculated note length ( = end time – start time) and note slope (= bandwidth/note length) before computing the final complexity metrics.

To capture song complexity, we first considered note count and note type, assuming that larger numbers of elements and unique elements provide the foundation of higher temporal and spectral complexity, respectively. To quantify spectral complexity, we calculated the standard deviations of note peak frequency, bandwidth and note slope. In the main analyses, we also z-standardised those three variables and summed up as a single metric (‘frequency variation’) to represent the overall spectral complexity of a song. To quantify temporal complexity, we calculated the standard deviation of note length, as well as song duration (= the end time of the last note – begin time of the first note) and note rate (= note count/song duration).

For single-note songs, we assigned zeros to the note variation-based metrics: frequency variation, standard deviations of peak frequency, bandwidth, and note length, to account for the absence of between-note variation. We also re-calculated note rate as the number of songs given per second (i.e., the total number of songs divided by the duration of all songs in the recording), to represent singing effect that is conceptually equivalent to the note rate measurement of multi-note songs. All complexity metrics for silent species (n = 12 species) were assigned zero to represent the lowest complexity.

### Sociality data

To quantify the level of sociality, we estimated the social group size of each species. A social group was defined as a stable aggregation of individuals that typically demonstrate strong social cohesion and interaction on a year-round basis. In classic group-living birds, group members spend most of the time together foraging, defending territory, bathing, roosting, keeping in contact with each other, warning each other of predators, and so on. The social group by definition reflects this consistent social unit that remains stable and interactive throughout the year. We exclude from this definition any temporary assemblies such as seasonal family groups that are observed after breeding, except in some socially monogamous tropical suboscines with high natal philopatry where offspring tend to remain for many months on the natal territory, foraging as a group with their parents [10]. We also exclude flocking behaviours associated with foraging during the non-breeding season, migratory journeys, or nocturnal roosts. The difference between these cases is that social groups are thought to develop a more complex and nuanced communication system (e.g. *Psophia* trumpeters, [11]), whereas other temporary aggregations are usually anti-predator strategies lacking a wider array of communication signals. We also exclude polygynous species that gather in single-sex groups at leks or display arenas. Although these aggregations may be stable over prolonged periods in some species (e.g. *Chiroxiphia* manakins), they do not fit our definition of social groups because they are purely based on mating displays, while the individuals concerned tend to be solitary in all other contexts away from the lek or display arena.

For each species, we collected data on the maximum social group size from species accounts in *Birds of the World* [3]. We focused on the maximum group size since it is likely the most robust estimate across a large number of species. Other metrics such as mean group size may more accurately reflect the average level of social interactions experienced by individuals. However, such estimates can only be derived from detailed population-level data and long-term monitoring studies, which are lacking for most suboscine species. Most information on social behaviour was found in sections on Behaviour and Breeding, but we also checked section on vocal behaviour and foraging behaviour for additional information [3]. In some cases, we derived maximum group sizes from the foraging behaviour of the species, on the basis that social species typically forage together and that foraging forms a major part of daily activities. We note that foraging descriptions may not always reflect consistent social behaviour, particularly for species that are difficult to survey or mostly encountered during breeding seasons only. We therefore adjusted scores in the context of other known aspects of life history such as migration, territoriality and social bond stability [12], which we assessed on a case-by-case basis. For example, migratory species generally do not maintain social bonds during migration or in the non-breeding season, so their group sizes were mainly assigned as one with rare exceptions.

### Phylogenetic modelling

#### Model specification

We have used several complexity metrics derived from standard deviations (SD) – frequency variation, and the SD of peak frequency, bandwidth, note slope and note length – to represent spectral and temporal variation. We assume that larger variation among notes will result in larger numerical values of SD. However, larger SDs may not necessarily translate into higher complexity as larger numbers inherently have larger SDs. For example, for a three-note song, the SD of three notes at the frequencies of 5-10-5 (kHz) is five times greater than three notes at frequencies of 1-2-1 (kHz), while the first song is only higher-pitched and not necessarily much more complex than the later. This mathematical issue may therefore inflate our SD-based complexity metrics for songs with higher frequencies and longer notes. SD-based metrics are more biologically meaningful when the values are compared against the same baseline, i.e., the same peak frequency for spectral metrics, and the same note length for the temporal metric. Therefore, in phylogenetic models, we added the mean peak frequency as an extra explanatory variable to models in which the response variable was frequency variation, peak frequency SD, bandwidth SD and note slope SD. Similarly, we added mean note length to the model where the response variable was note length SD.

## Supplementary figures

**Figure S1.**
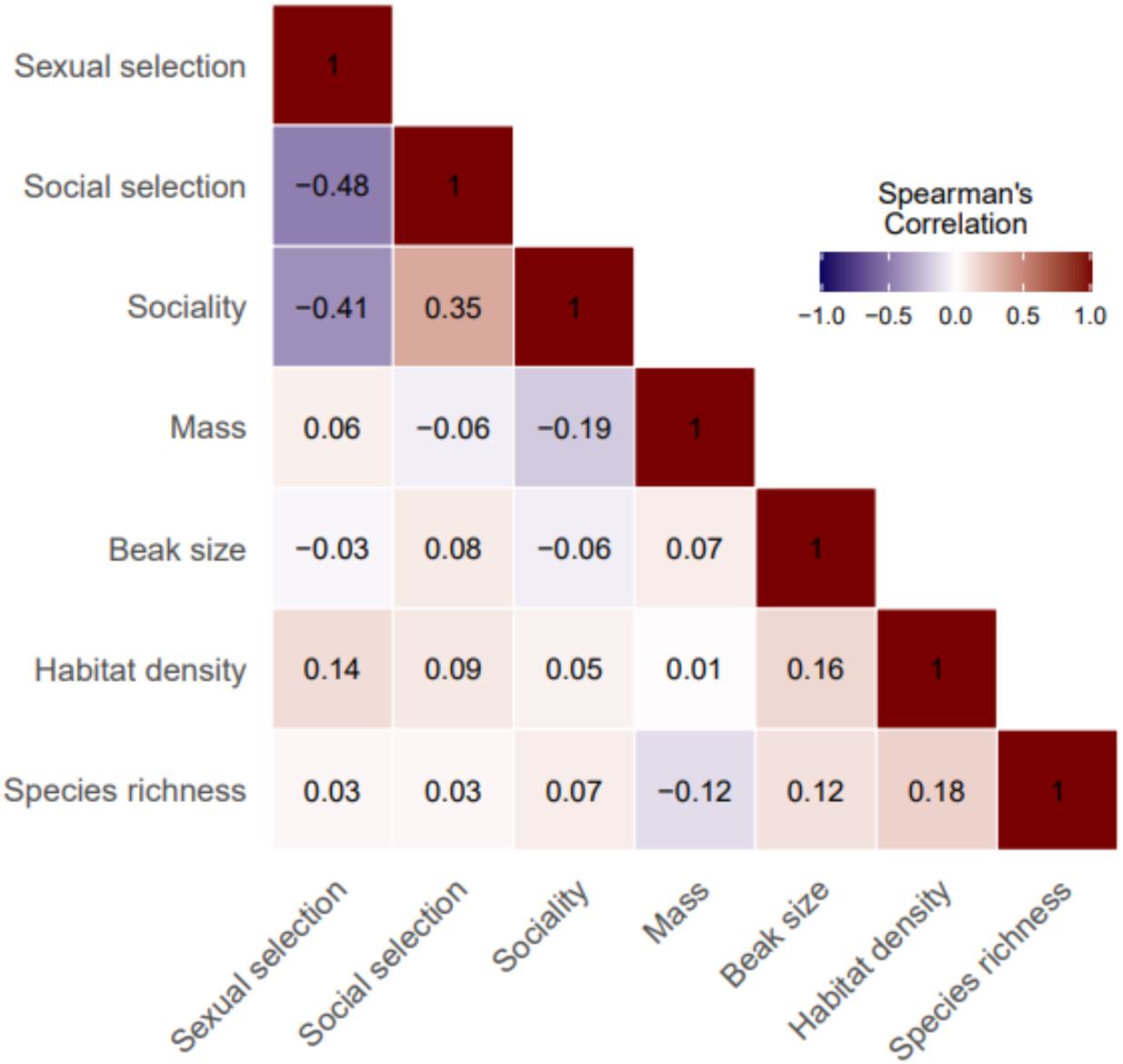
Correlation between main explanatory variables. Correlation plot showing Spearman’s correlation coefficients between the tested explanatory variables. Results are based on data from all suboscine passerine bird species (n = 1,276 species). Warmer colours indicate stronger positive correlation; cooler colours indicate stronger negative correlation. See Methods and Data S1A for detailed definitions of each variable.

**Figure S2.**
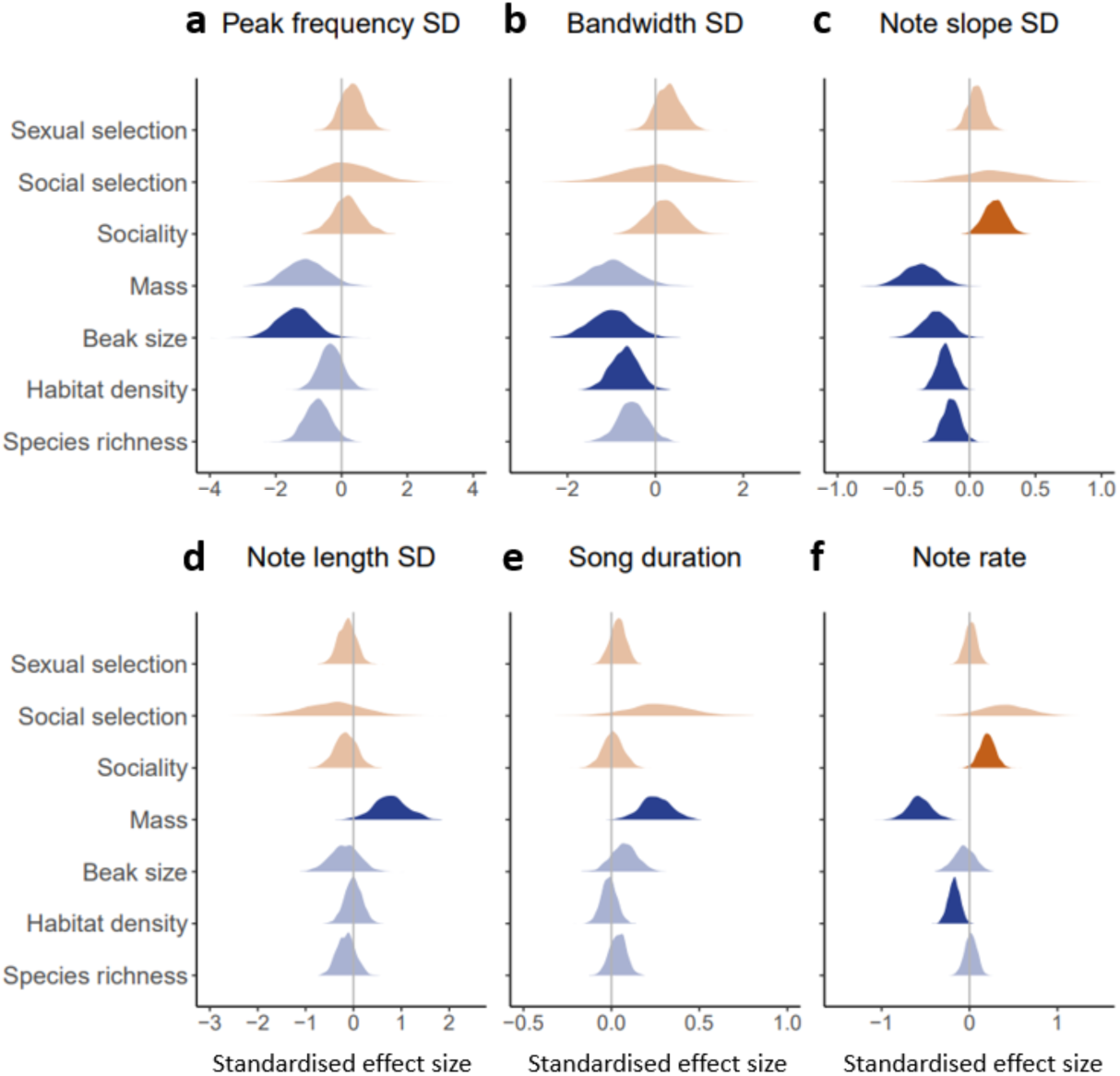
Results of Bayesian phylogenetic models using additional complexity metrics. Panels show the results of Bayesian phylogenetic models testing the effects of the proposed social drivers (orange) and ecological constraints (blue) on the evolution of song complexity in 1,276 species of suboscine passerine birds. Spectral complexity was measured as the standard deviation of peak frequency, bandwidth and note slope (a-c). Temporal complexity was measured as the standard deviation of note length, song duration and note rate (d-f). Models used the phylogenetic tree provided in Harvey et al. [2]. To account for the statistical artefact related to complexity metrics using standard deviations (see electronic supplementary material), we added the average note frequency to spectral complexity models(a-c), and average note length to temporal complexity models (d). These additional explanatory variables are not shown in this figure to reduce clutter and improve visual clarity. Shaded areas show the distribution of estimated effect sizes of the predictor, based on a sample of 2,000 posterior draws. Dark shades indicate statistical significance where 95% credible intervals do not overlap with zero.

## Supplementary tables

**Table S1.** Definition of song complexity metrics used in this study. Metrics denoted with an asterisk (*) are used in the main analyses. SD: standard deviation. For one-note songs, the metrics based on standard deviations (frequency variation and metrics ending with SD) were assigned zero to reflect the absence of note variation within a song. For silent species, all metrics were assigned zero to reflect the lowest signal complexity.

| Complexity metric | Description |
| --- | --- |
| Note count * | The total number of notes present in the song |
| Number of note types * | The number of unique note types present in the song |
| Peak frequency SD | The standard deviation of peak frequency of all notes in the song. Peak frequency (Hz) is the frequency where the note reaches its highest amplitude |
| Bandwidth SD | The standard deviation of bandwidth of all notes in the song. Bandwidth (Hz) is the difference between the two frequency limits spanning 5% and 95% of the energy of each note |
| Note slope SD | The standard deviation of note slope of all notes in the song. Note slope indicates the flatness of the note shape, calculated as bandwidth/note length |
| Frequency variation * | The sum of z-standardised peak frequency SD, bandwidth SD and note slope SD of the song |
| Note length SD | The standard deviation of note length of all notes in the song. Note length (ms) is the duration between the begin time and end time of the note |
| Song duration | The duration (s) between the begin time of the first note and the end time of the last note. For one-note songs, song duration equals to the note length |
| Note rate | The number of notes given per second. For multi-note songs, note rate = note count / song duration; for one-note songs, note rate = the number of songs per second, calculated as the number of songs divided by the duration of all songs present in the recording |

**Table S2.**
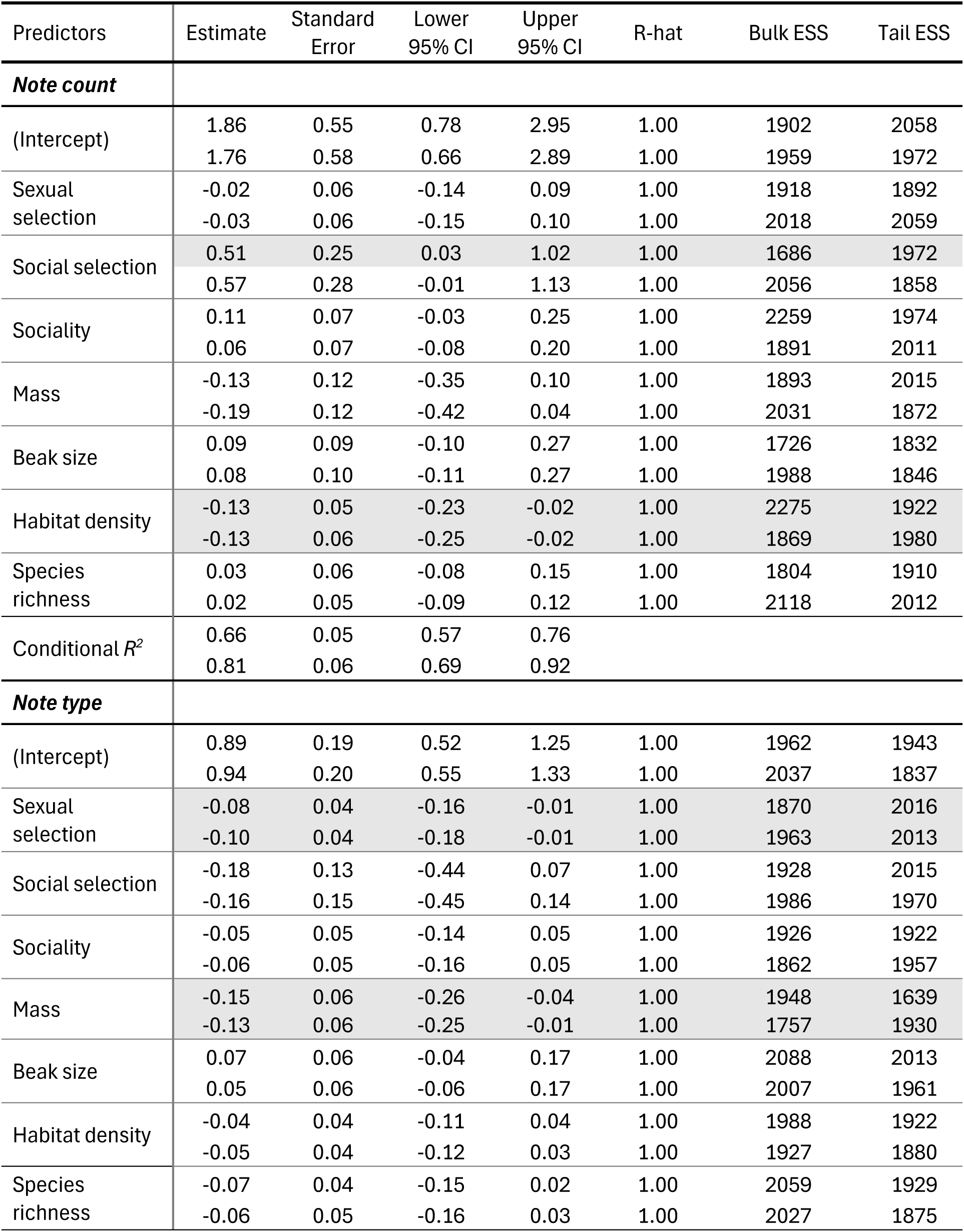
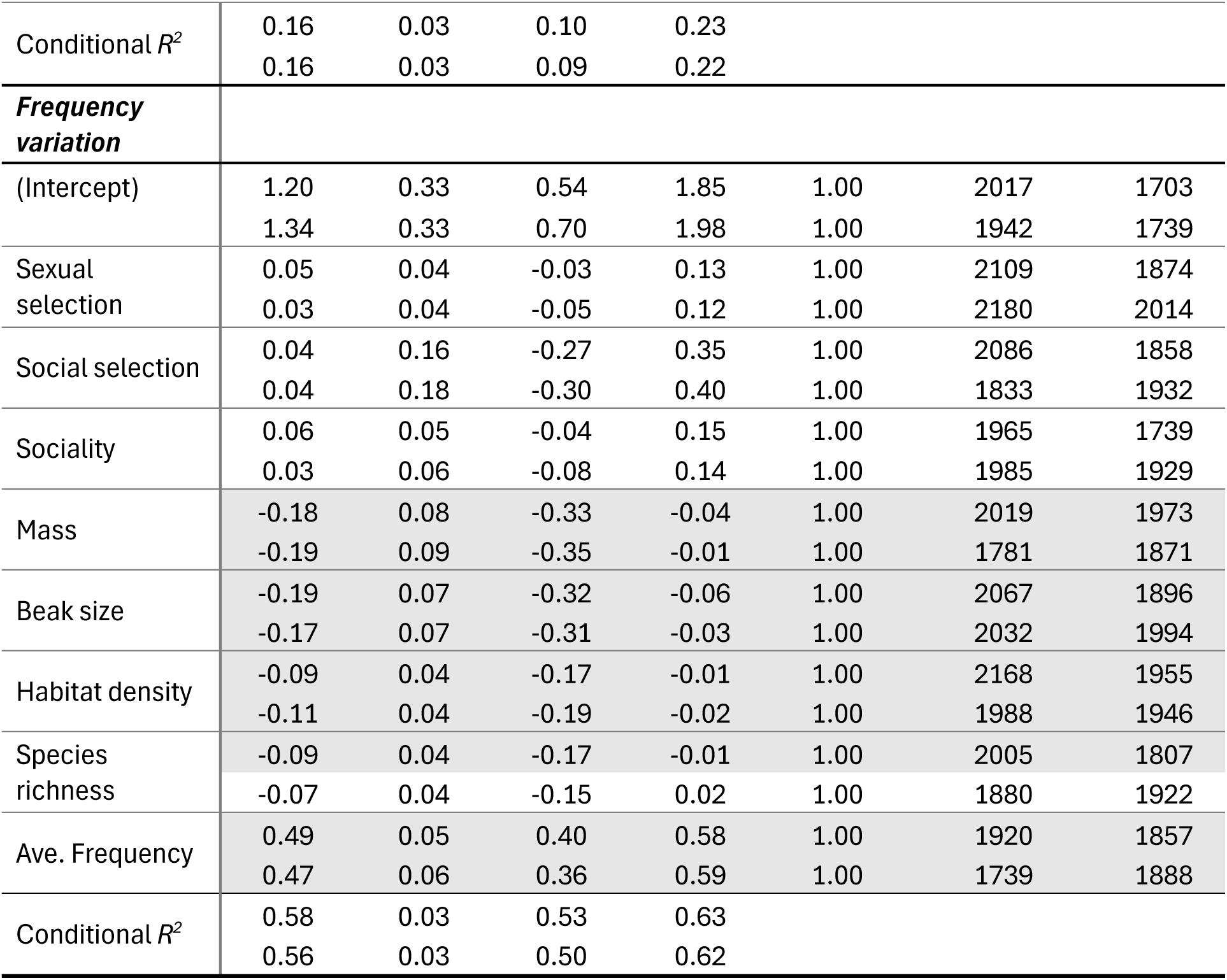
Results of Bayesian phylogenetic models. For each predictor and conditional *R^2^*, we show results using all species (N = 1,276 species; top row) and species with high-certainty data (N = 1,028 species; bottom row). Predictors showing significant effects (95% CI not spanning zero) on metrics of complexity are highlighted in grey. All results of each model were based on a sample of 2,000 posterior draws (see Methods). CI: credible interval; ESS: effective sample size. An R-hat value approximating one, and ESS values greater than 1,000 indicate that the model was converged and could accurately estimate the posterior distribution of the parameter [13].

| Predictors | Estimate | Standard Error | Lower 95% CI | Upper 95% CI | R-hat | Bulk ESS | Tail ESS |
| --- | --- | --- | --- | --- | --- | --- | --- |
| **Note count** |  |  |  |  |  |  |  |
| (Intercept) | 1.86 | 0.55 | 0.78 | 2.95 | 1.00 | 1902 | 2058 |
|  | 1.76 | 0.58 | 0.66 | 2.89 | 1.00 | 1959 | 1972 |
| Sexual selection | -0.02 | 0.06 | -0.14 | 0.09 | 1.00 | 1918 | 1892 |
|  | -0.03 | 0.06 | -0.15 | 0.10 | 1.00 | 2018 | 2059 |
| Social selection | 0.51 | 0.25 | 0.03 | 1.02 | 1.00 | 1686 | 1972 |
|  | 0.57 | 0.28 | -0.01 | 1.13 | 1.00 | 2056 | 1858 |
| Sociality | 0.11 | 0.07 | -0.03 | 0.25 | 1.00 | 2259 | 1974 |
|  | 0.06 | 0.07 | -0.08 | 0.20 | 1.00 | 1891 | 2011 |
| Mass | -0.13 | 0.12 | -0.35 | 0.10 | 1.00 | 1893 | 2015 |
|  | -0.19 | 0.12 | -0.42 | 0.04 | 1.00 | 2031 | 1872 |
| Beak size | 0.09 | 0.09 | -0.10 | 0.27 | 1.00 | 1726 | 1832 |
|  | 0.08 | 0.10 | -0.11 | 0.27 | 1.00 | 1988 | 1846 |
| Habitat density | -0.13 | 0.05 | -0.23 | -0.02 | 1.00 | 2275 | 1922 |
|  | -0.13 | 0.06 | -0.25 | -0.02 | 1.00 | 1869 | 1980 |
| Species richness | 0.03 | 0.06 | -0.08 | 0.15 | 1.00 | 1804 | 1910 |
|  | 0.02 | 0.05 | -0.09 | 0.12 | 1.00 | 2118 | 2012 |
| Conditional $R^2$ | 0.66 | 0.05 | 0.57 | 0.76 | | | |
|  | 0.81 | 0.06 | 0.69 | 0.92 |  |  |  |
| **Note type** |  |  |  |  |  |  |  |
| (Intercept) | 0.89 | 0.19 | 0.52 | 1.25 | 1.00 | 1962 | 1943 |
|  | 0.94 | 0.20 | 0.55 | 1.33 | 1.00 | 2037 | 1837 |
| Sexual selection | -0.08 | 0.04 | -0.16 | -0.01 | 1.00 | 1870 | 2016 |
|  | -0.10 | 0.04 | -0.18 | -0.01 | 1.00 | 1963 | 2013 |
| Social selection | -0.18 | 0.13 | -0.44 | 0.07 | 1.00 | 1928 | 2015 |
|  | -0.16 | 0.15 | -0.45 | 0.14 | 1.00 | 1986 | 1970 |
| Sociality | -0.05 | 0.05 | -0.14 | 0.05 | 1.00 | 1926 | 1922 |
|  | -0.06 | 0.05 | -0.16 | 0.05 | 1.00 | 1862 | 1957 |
| Mass | -0.15 | 0.06 | -0.26 | -0.04 | 1.00 | 1948 | 1639 |
|  | -0.13 | 0.06 | -0.25 | -0.01 | 1.00 | 1757 | 1930 |
| Beak size | 0.07 | 0.06 | -0.04 | 0.17 | 1.00 | 2088 | 2013 |
|  | 0.05 | 0.06 | -0.06 | 0.17 | 1.00 | 2007 | 1961 |
| Habitat density | -0.04 | 0.04 | -0.11 | 0.04 | 1.00 | 1988 | 1922 |
|  | -0.05 | 0.04 | -0.12 | 0.03 | 1.00 | 1927 | 1880 |
| Species richness | -0.07 | 0.04 | -0.15 | 0.02 | 1.00 | 2059 | 1929 |
|  | -0.06 | 0.05 | -0.16 | 0.03 | 1.00 | 2027 | 1875 |
| Conditional $R^2$ | 0.16 | 0.03 | 0.10 | 0.23 | | | |
|  | 0.16 | 0.03 | 0.09 | 0.22 |  |  |  |
| **Frequency variation** |  |  |  |  |  |  |  |
| (Intercept) | 1.20 | 0.33 | 0.54 | 1.85 | 1.00 | 2017 | 1703 |
|  | 1.34 | 0.33 | 0.70 | 1.98 | 1.00 | 1942 | 1739 |
| Sexual selection | 0.05 | 0.04 | -0.03 | 0.13 | 1.00 | 2109 | 1874 |
|  | 0.03 | 0.04 | -0.05 | 0.12 | 1.00 | 2180 | 2014 |
| Social selection | 0.04 | 0.16 | -0.27 | 0.35 | 1.00 | 2086 | 1858 |
|  | 0.04 | 0.18 | -0.30 | 0.40 | 1.00 | 1833 | 1932 |
| Sociality | 0.06 | 0.05 | -0.04 | 0.15 | 1.00 | 1965 | 1739 |
|  | 0.03 | 0.06 | -0.08 | 0.14 | 1.00 | 1985 | 1929 |
| Mass | -0.18 | 0.08 | -0.33 | -0.04 | 1.00 | 2019 | 1973 |
|  | -0.19 | 0.09 | -0.35 | -0.01 | 1.00 | 1781 | 1871 |
| Beak size | -0.19 | 0.07 | -0.32 | -0.06 | 1.00 | 2067 | 1896 |
|  | -0.17 | 0.07 | -0.31 | -0.03 | 1.00 | 2032 | 1994 |
| Habitat density | -0.09 | 0.04 | -0.17 | -0.01 | 1.00 | 2168 | 1955 |
|  | -0.11 | 0.04 | -0.19 | -0.02 | 1.00 | 1988 | 1946 |
| Species richness | -0.09 | 0.04 | -0.17 | -0.01 | 1.00 | 2005 | 1807 |
|  | -0.07 | 0.04 | -0.15 | 0.02 | 1.00 | 1880 | 1922 |
| Ave. Frequency | 0.49 | 0.05 | 0.40 | 0.58 | 1.00 | 1920 | 1857 |
|  | 0.47 | 0.06 | 0.36 | 0.59 | 1.00 | 1739 | 1888 |
| Conditional $R^2$ | 0.58 | 0.03 | 0.53 | 0.63 | | | |
|  | 0.56 | 0.03 | 0.50 | 0.62 |  |  |  |

